# Jet-printing microfluidic devices on demand

**DOI:** 10.1101/2020.05.31.126300

**Authors:** Cristian Soitu, Nicholas Stovall-Kurtz, Cyril Deroy, Alfonso A. Castrejón-Pita, Peter R. Cook, Edmond J. Walsh

## Abstract

There is an unmet demand for microfluidics in biomedicine. We describe contactless fabrication of microfluidic circuits on standard Petri dishes using just a dispensing needle, syringe pump, 3-way traverse, cell-culture media, and an immiscible fluorocarbon (FC40). A submerged micro-jet of FC40 is projected through FC40 and media on to the bottom of a dish, where it washes media away to leave liquid fluorocarbon walls pinned to the substrate by interfacial forces. Such fluid walls can be built into almost any imaginable 2D circuit in minutes, which we exploit to clone cells using limiting dilution in a way that beats the Poisson limit, sub-culture adherent cells, and feed arrays of cells continuously for a week. This general method should have wide application in biomedicine.

**One sentence summary:** In the everyday world, we cannot build complex structures out of liquids as they collapse into puddles; in the microworld we can.

## INTRODUCTION

As sensitivities of methods for detecting biomolecules improve, demand for handling ever-smaller volumes increases – and this drives development of microfluidic approaches (Sackmann *et al*., 2014; Prakadan *et al*., 2017; Sonnen and Merten, 2019). However, few of these are found in biomedical workflows, with the exception of those involving microplates. Why? Reasons given include: devices are expensive and take days/weeks to make, they are complicated to operate, their contents are inaccessible, and they are not made with cell-friendly materials (Sackmann *et al*., 2014).

In the everyday world, gravity is such a dominant force that most objects are made with solids, and one cannot contemplate building them out of liquids, which just collapse into puddles. Consequently, liquids are always contained by solid walls, otherwise they drain away. In the microworld, gravity becomes irrelevant, and interfacial forces dominate (think of water-striders skimming over ponds, and dew-drops sticking to blades of grass). Consequently, an interface between two fluids can act as a robust wall separating the two. This paradigm enabled emergence of open microfluidics (Berthier *et al.*, 2016), where solid walls are replaced by air:water (Popova *et al.*, 2015) or oil:water interfaces (Walsh *et al*., 2017; Soitu *et al*., 2018; Chao *et al.*, 2020).

Open microfluidic devices are generally easier to integrate into biomedical workflows as they offer better optical and physical access to samples, reduce adhesion of reagents to solid surfaces and are resistant to air bubbles. However, despite the ever-increasing efforts to simplify manufacturing workflows for such devices, most of them still require etching of the substrate (Berthier *et al.*, 2019), surface treatment (Li *at al.*,2016) or contact (Soitu *et al*., 2018), or some combination of these (Chao *et al.*, 2020) to confine fluidic structures. Such complex manufacturing processes deter many biologists who favor fast flexible prototyping without having to compromise biocompatibility (Berthier *et al.*, 2012).

Here, we describe a contactless method to fabricate microfluidic devices on demand, where the only ‘building’ materials used are those in the biocompatible trio – cell media, the immiscible fluorocarbon FC40, and a polystyrene Petri dish. FC40 is ‘jetted’ from a dispensing needle through bulk FC40 and media on to the untreated bottom of the dish. Complex microfluidic structures can be produced reproducibly with high accuracy (e.g., with features <50 μm in size) in minutes. The aqueous phase is confined by fluid walls – media:FC40 interfaces – which are robust yet easily pierced (so liquids can be added/removed through them at any preselected point) whilst being transparent. The physics underlying flow during such jetting is complex (Glauert, 1956; Deshpande and Vaishnav, 1982; Phares *et al.*, 2000; Davis *et al*., 2012); therefore, we establish appropriate conditions. We then exploit jetting to ‘beat’ the Poisson limit to clone single mammalian cells by limited dilution, sub-culture them (again using a contactless method), and perfuse them steadily with fresh media for a week.

## RESULTS

### Approach

**Figure 1A** illustrates the approach. The bottom of a standard tissue-culture dish is covered with a film of cell-growth media, and an FC40 overlay added to prevent evaporation. A dispensing needle filled with FC40, connected to a syringe pump, and held by a 3-way traverse is now lowered below the surface of the fluorocarbon; starting the pump jets FC40 on to the dish to push media aside. As FC40 has a low equilibrium contact angle (CA) on polystyrene (< 10°), it wets it better than media (equilibrium CA ∼50°; Walsh *et al.*, 2017), so it adheres to the bottom. Moving the micro-jet sideways then creates a line of FC40 on the dish, and drawing more lines creates a grid with 256 chambers in <2 min (**Fig. 1B**; **Movie 1**). Each chamber is isolated from others by liquid walls of FC40 pinned to polystyrene. Interfacial forces dictate chamber geometry – a spherical cap sitting on a square footprint (height ∼75 µm; volume ∼100 nl). Up to ∼900 nl more media can be pipetted into chambers as fluid walls morph above unchanging footprints. Chambers are then used like wells in microplates: liquids are added/removed to/from them by pipetting through FC40 instead of air. The maximum and minimum volumes that can be held in chambers without altering footprints are determined by advancing and receding contact angles; addition of too much media inevitably merges adjacent chambers. Even so, chambers accept a manyfold wider range of volume than equally-spaced wells in a microplate, whilst containing ∼1,000^th^ the volume (Soitu *et al*., 2018). Consequently, if chambers contain cells, the volume ratio of intra- to extra-cellular fluid more closely resembles that *in vivo*. Importantly, this method is contactless: the nozzle touches neither dish nor media. Moreover, one pipet tip can add/reagents to/from many chambers without detectable cross-contamination (shown – for example – by seeding bacteria in every other chamber, adding media to all through one tip, and finding that bacteria grow only in inoculated chambers as others remain sterile; Soitu *et al*., 2018). In other words, a tip is washed effectively by passage through FC40 between chambers, and – when using cells – we make doubly sure by additionally washing in 70% ethanol.

**Figure 1.**
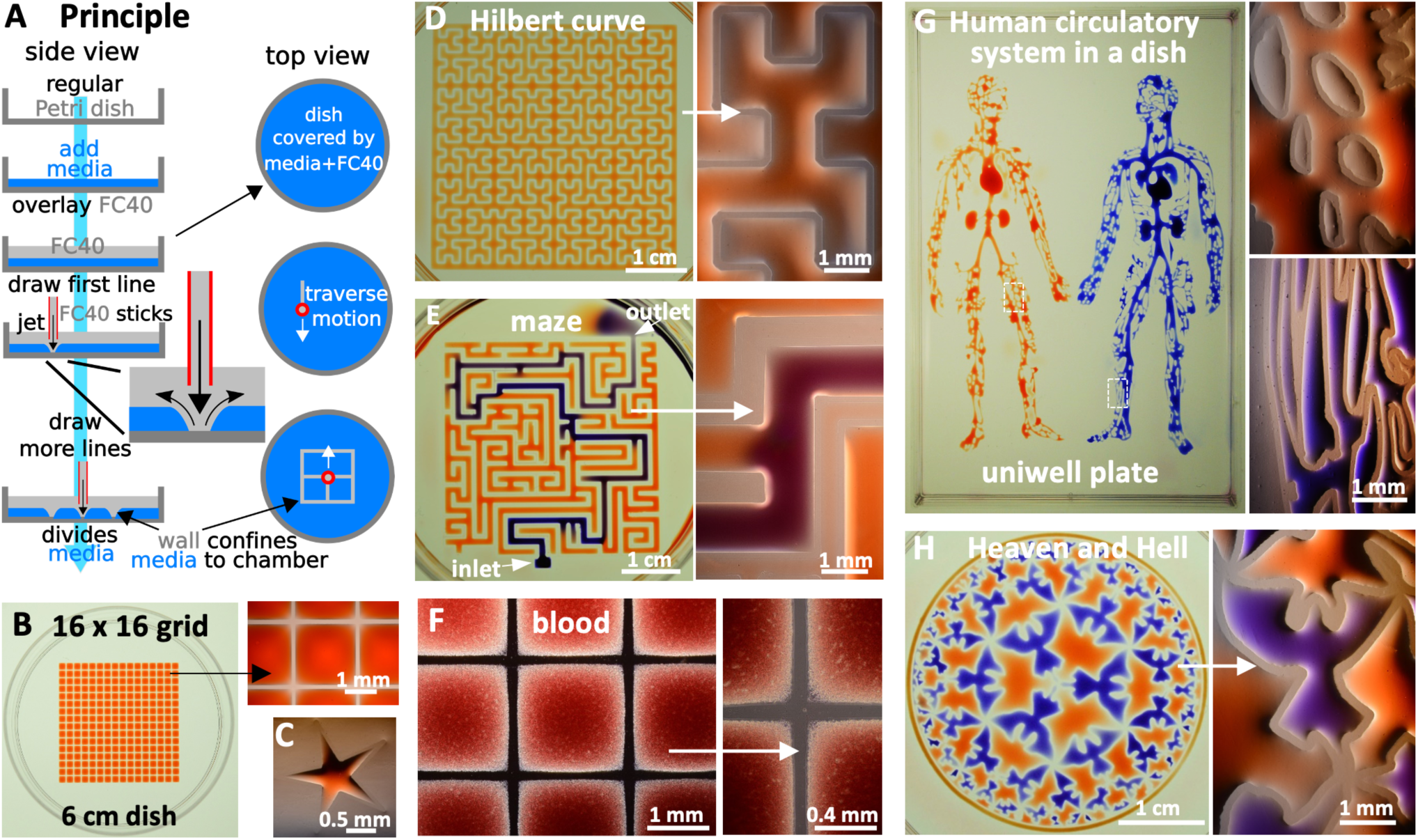
Principle. **A**. A ‘micro-jet’ of an immiscible fluorocarbon (FC40) is projected through FC40 and media on to the bottom of a dish; media is pushed aside, and FC40 sticks to the bottom (it wets polystyrene better than media). Moving the jet now draws lines to form a grid with FC40 walls. **B**. A 16 x 16 grid (inter-chamber spacing 1.9 mm; 600 nl red dye added to each chamber; zoom shows fluid walls). **C**. Star-shaped walls (red dye added after jetting). **D**. A square with one internal wall shaped like a Hilbert curve (a continuous line with many 90° and 45° turns). Red dye was added at many points; it diffused around the wall throughout the square. Zoom shows some turns. **E**. Path through a maze with a single inlet and outlet. The circuit was built by jetting FC40 through media plus red dye; when blue dye is pumped into the inlet, it takes the path of least resistance through the maze. Zoom shows some walls. **F**. Part of 16 x 16 grid where FC40 is jetted through blood instead of media. Zoom shows the jet leaves no cells in what are now FC40 walls. **G.** Human circulatory system in a uniwell plate with dimensions of a 96-well plate. Red and blue dyes were infused into major ‘arteries’ and ‘veins’ which then flow/diffuse throughout each system. Zooms show regions in white rectangles. **H**. Walls were built to reproduce the Circle Limit IV (‘Heaven and Hell’) by M.C. Escher – a circular, regular tiling, logically bounded on all sides by the infinitely small – and blue and red dyes added to ‘angels’ and ‘devils’ (zoom).

Jetting allows walls to be built with a precision only limited by that of the traverse – illustrated by building a star-shaped chamber (**Fig. 1C**), a square enclosing a single continuous wall shaped like a Hilbert curve (**Fig. 1D**; fabrication time ∼5 min), and a microfluidic maze where blue dye infused into the inlet solves the problem of finding the path to the sole exit (**Fig. 1E**; conduits have maximum widths of 1 mm, and heights of ∼350 µm). Reproducibility is demonstrated by constructing a fractal circuit in which a central input is connected through conduits (footprint width 450 µm, maximum height ∼160 µm) that each make 5 right-angled turns to one of 64 outlets; dye infused into the input reaches all outlets simultaneously, showing that conduit cross-sections and lengths are similar throughout (**Fig. S1**). The method is not restricted to media, which can be replaced by whole human blood (**Fig. 1F**). Nor is it restricted to regular patterns, illustrated by the ‘human circulatory system’ (**Fig 1G**; jetting time ∼90 min) and M.C. Escher’s ‘Heaven and Hell’ (**Fig. 1H**; jetting time ∼5 min). In these and subsequent examples, dyes are added solely to aid visualization; they play no role in stabilizing liquid structures. We also use ‘media’ to describe DMEM plus 10% FBS unless stated otherwise. In addition, fluid walls are sufficiently stable to be carried up/down stairs between labs, incubators, and microscopes – just like any dish filled with liquid (see **Movie 2**). These examples illustrate the versatility of jetting and show that circuits with almost any imaginable 2D shape can be built quickly and accurately using cell-friendly materials.

**Figure S1.**
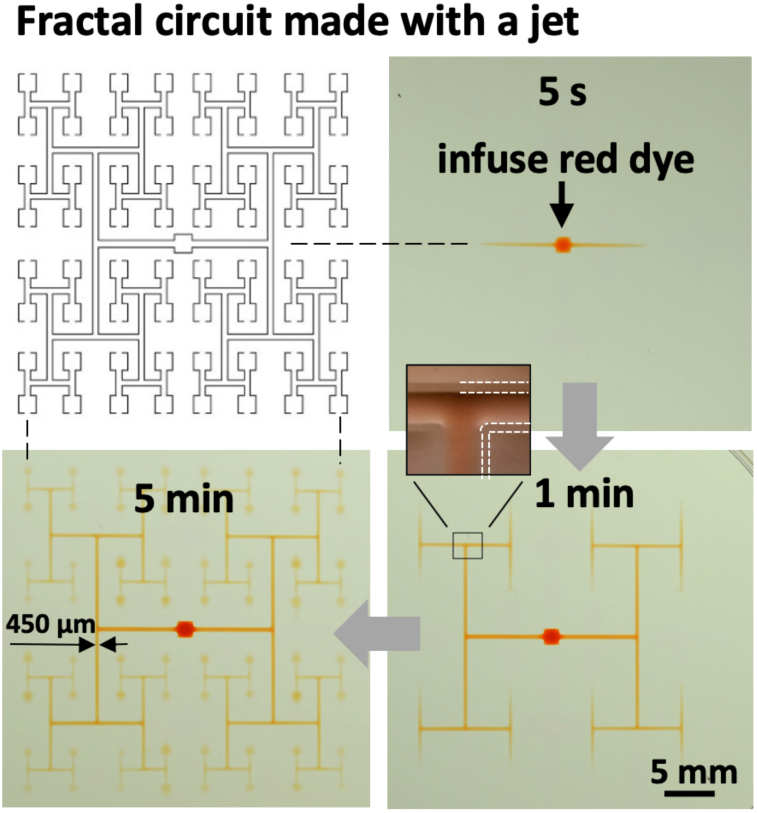
Making a fractal circuit using a jet. The jet provides great accuracy and precision, as the jetting needle is held by the traverse at a defined *x,y,z* position (so there is no play in the system), and it contacts neither media nor dish. This circuit is constructed with high accuracy in ∼6 min from T-shaped segments (plan shown at top left; conduits have footprint widths of 450 µm) connecting a central input to 64 outlets and the rest of the dish (the sink). Three images show the circuit in operation; red dye was infused into the input and it reaches each outlet simultaneously, showing that all branches have similar dimensions. Inset: one junction with some wall edges highlighted white.

### Building walls and conduits with different widths

In theory, many parameters including interfacial properties, jet momentum, densities, and kinematic viscosities influence wall stability and shape. FC40 has a density of 1.855 g/ml, and intuition suggests that buoyancy should force media above the fluorocarbon; however, media remains pinned to a pre-wetted dish as interfacial forces are stronger than buoyancy forces. Other parameters were varied to assess their impact on wall construction (**Fig. 2A**; **Fig. S2A**). When the nozzle is high above the substrate (i.e., *H* is large), no walls form; the momentum of the jet falls as it travels through FC40 (which has a kinematic viscosity of 2.2 cSt compared to 1 for water), making it insufficient to displace media (**Fig. 2Bi**, red symbol). Lowering the nozzle decreases the loss to the point when the final jet momentum that hits the substrate is large enough to push media aside. FC40 then remains pinned to the substrate to give a wall. Examination of streamlines (**Movie 3**) shows that jet width increases with distance from the nozzle, so further lowering yields progressively narrower walls as the jet impinges on progressively-smaller substrate areas. Even so, wall width proves to be relatively tolerant to changes in *H* (**Fig. 2Bi**), so walls can be constructed using a nozzle positioned at the same height above dishes from different manufacturers that turn out to be slightly bowed to different degrees. At a low volumetric flow rate 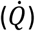, momentum is insufficient and no wall forms; increasing 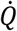 yields walls with increasing width **(Fig. 2Bii)**. Nozzle diameter (*D*_*nozzle*_), traverse velocity (*V*_*traverse*_) and interfacial properties (especially presence of FBS) all affect wall width (**Fig. 2Bii,iii**; see **Fig. S2B** for data on E8+, mTesR, StemFlex media). Wider walls can be made by overlapping jetting lines (**Fig. 2C**). Conduit width is easily adjusted down to at least 35 µm by jetting walls closer together (**Fig. 2D**). We have therefore identified a range of conditions allowing wall formation even on commercially-available dishes that are not completely flat.

**Figure 2.**
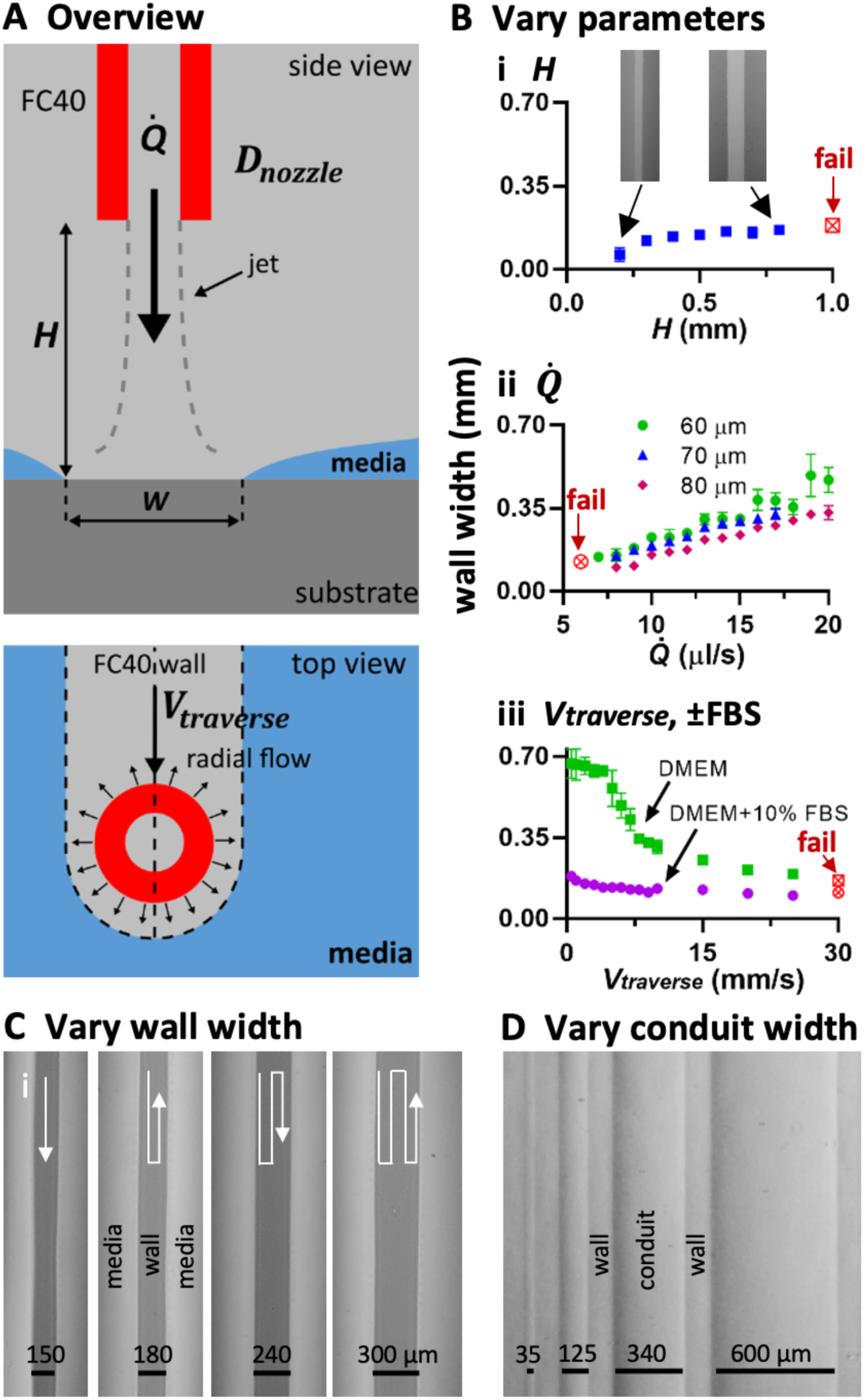
Varying jetting parameters (*W*). Unless stated otherwise throughout the paper, 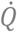 (volumetric flow rate) = 8 μl/s, *H* (height of nozzle above substrate) = 0.4 mm, *D*_*nozzle*_ (diameter of jetting nozzle) = 60 µm, *V*_*traverse*_ (lateral traverse speed) = 10 mm/s, and media was DMEM + 10% FBS. **A.** Overview. A submerged FC40 micro-jet drives media off the substrate to create an FC40 wall, and some parameters varied are indicated. **B.** Controlling wall width (red symbols: continuous walls fail to form) by varying: (**i)** jet height (inset: images of walls when *H* = 0.2 and 0.8 mm). As the nozzle is lowered closer to the substrate, less jet momentum is lost to the bulk FC40, and the net momentum hitting the substrate increases, resulting in wider walls. (**ii**) flow rate and nozzle diameter. Wall width increases as jet momentum increases (by increasing flowrate or reducing nozzle diameter) (**iii**) traverse velocity and serum content. Surface tension between media and the substrate increases with serum content, so the ‘sweeping’ action of the jet momentum is diminished, resulting in smaller wall widths. **C.** Varying wall width by overlapping walls. Wall width can be increased by setting the separation between jetting paths to be smaller than the wall width generated by one pass. **D.** Making conduits with different widths (walls all have equal widths) by varying the distance between consecutive paths. *H*= 0.3 mm, 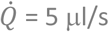, *D*_*nozzle*_ = 60 µm, *V*_*traverse*_ = 1.8 mm/s.

**Figure S2.**
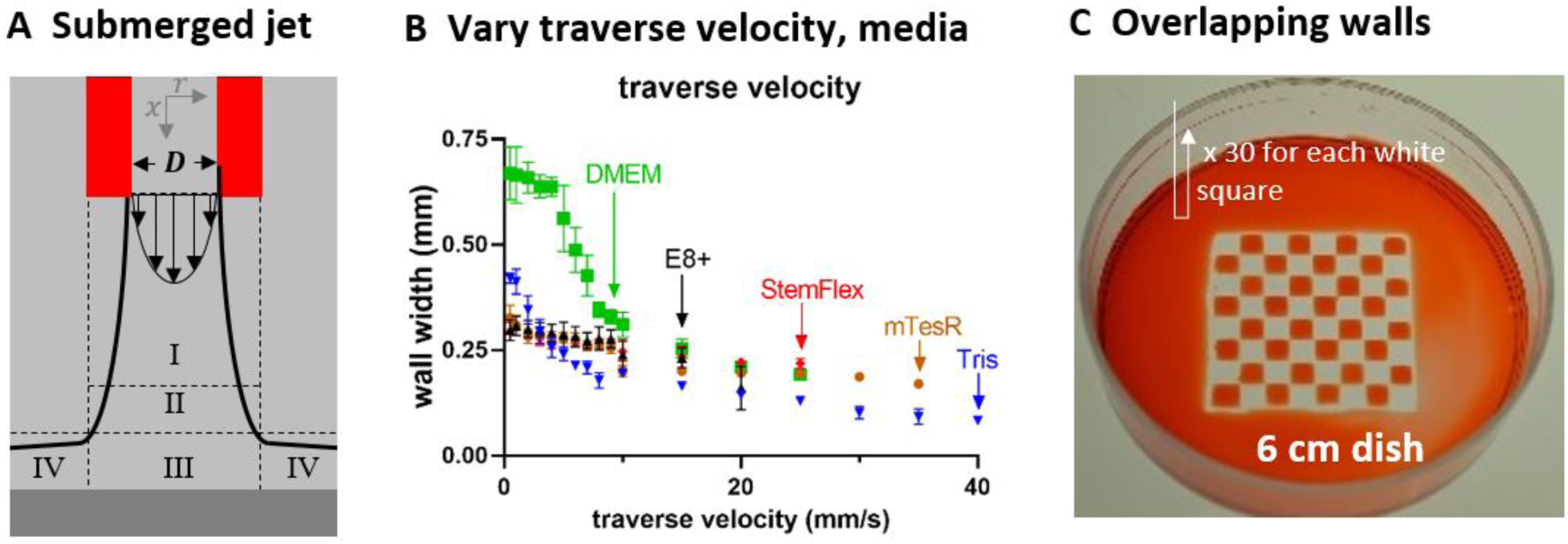
**A.** Flow profiles in/around a submerged jet impinging on a flat substrate (Deshpande and Vaishnav, 1982). Five arrows illustrate the flow profile on emergence from the nozzle, and then there are 4 regions of jet flow assuming a fully-developed exit-flow velocity-profile. I: free jet region where jet diameter progressively increases. II: transition-to-wall zone. III: stagnation zone. IV: wall-jet region. In our case, the substrate is initially covered with media and removed by jet momentum and the shearing force in the wall-jet zone as the jet traverses. **B.** Effect of traverse velocity on wall width using different media (*H* = 0.5 mm, 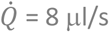; 4 mg/ml resazurin dye added to 20 mM TrisHCl). No media tested gave stable walls with *V*_*traverse*_ >35 mm/s (as dwell time over any one area in the dish is too short for the jet to remove media from the surface). **C.** Building wide overlapping walls (3 x 3 mm) by jetting through media + dye.

### Controlling flow through circuits

Flow through these circuits can easily be driven by external pumps; fluid walls/ceilings spontaneously seal around connecting hydrophilic tubes on insertion, and then morph above unchanging footprints during flow. However, adding external pumps adds complexity and cost. As intrinsic differences in Laplace pressure can drive flow through microfluidic circuits (Walker and Beebe, 2002; Walsh *et al*., 2017), we demonstrated this in these circuits (**Fig. S3)**.

Thus far, we have jetted FC40. We now create a microfluidic ‘valve’ by jetting media (**Fig. S4, Movie 4**). We first build a conduit with a dead-end (using an FC40 jet), and make a hole through the end wall (using a media jet traversing down the conduit axis) to allow flow through the hole. We next rebuild the original end wall (using an FC40 jet that traverses across the conduit) to stop flow. Such valves can be opened and closed as many times as needed.

**Figure S3.**
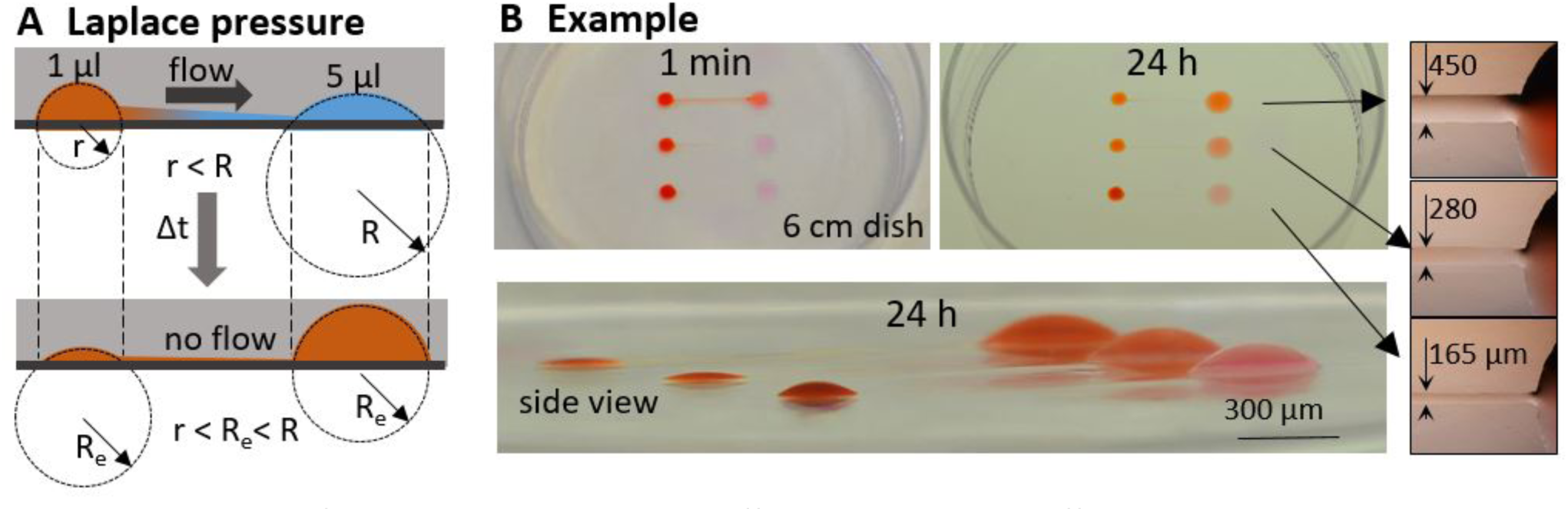
Controlling flow through conduits with different widths using differences in Laplace pressure. **A.** Laplace pressure – principle. Top: a difference in Laplace pressure spontaneously drives flow as drops have different radii of curvature (r<R). Bottom; flow ceases when both drops have the same curvature and radius at equilibrium, R_e_. **B.** Example. Each of the 3 circuits was made by jetting one continuous wall; all 3 are initially identical (left- and right-hand chambers have diameters of 2.5 and 5 mm respectively) except for conduit width which varies from 450-165 µm. Red dye (5 µl) is added to left-hand drops to increase Laplace pressure to drive flow to the right. After 1 min, some dye in the upper circuit (but not others) reaches the right-hand drop. After 24 h, the left-hand upper drop is nearly flat (and so empty) as essentially all dye has been discharged through the widest conduit to the right-hand one. In contrast, the left-hand lower drop remains rounded and continues to drive flow through the narrowest conduit. Insets: zooms of conduits with widths indicated.

**Figure S4.**
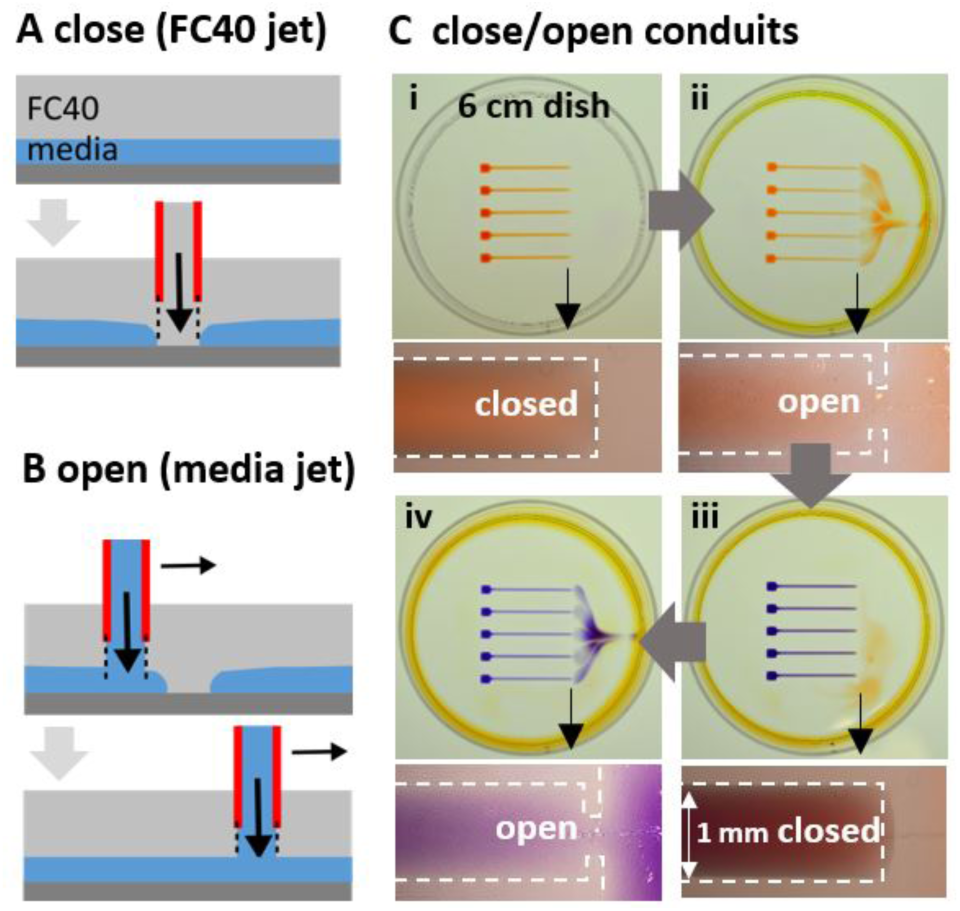
A microfluidic valve. **A.** Closing an open conduit. A new wall across a conduit is built by jetting FC40 (nozzle moves into page). **B.** Opening a closed conduit. A wall across a conduit is removed by jetting media through the wall (nozzle moves to right). **C.** Example. Insets show conduit ends, with dashed lines marking FC40 walls. (**i**). Five continuous FC40 walls were printed to create 5 identical circuits (each a reservoir with a 2 x 2 mm footprint connected to a conduit 1 mm wide), and 2 µl red dye added to each reservoir. Red dye is confined to each circuit as the surrounding FC40 wall is continuous. (**ii**) Media was jetted (*H* = 0.4 mm, 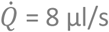, *V*_*traverse*_ *=* 15 mm/s) to open end walls. Red dye spills out into the dish (which acts as a sink, as it has a low Laplace pressure). (**iii**) Ends of conduits were closed by jetting FC40 (conditions as for media) to rebuild walls, and 2 µl blue dye added to reservoirs to test for leaks. No blue dye escapes, showing new walls provide perfect seals. (**iv**) Reopening end walls allows escape of blue dye. The pattern indicates flow through each end is similar, so openings have equal widths.

### ‘Beating’ the Poisson limit during cell cloning

Mammalian cells are often cloned by splitting a dilute cell suspension amongst wells in a microplate so most wells get no cells, a few get one, and fewer still get two or more – before singletons are selected, grown, and resulting colonies picked. This approach is wasteful, as Poisson statistics ensure that wells with singletons are in a minority. Although many approaches (including fluorescence activated cell sorting, and combinations of printing plus imaging; Yim and Shaw, 2018) can increase the number of wells with singletons and so ‘beat’ the Poisson limit, cost and complexity prevent widespread adoption (Collins *et al*., 2015; Welch and Arden, 2019). Moreover, solid plastic walls around each well in a microplate yield ‘edge effects’ that obscure cells close to walls (Soitu *et al*., 2019b). Such effects are so severe that many users never check to see whether a well contains just one cell immediately after plating, and go through a 2^nd^ cloning round to increase the chances of achieving monoclonality. This prompted development of an approach that allies jetting with use of Voronoi diagrams.

The approach is illustrated using 100 randomly-distributed dots as cell surrogates printed on a transparency stuck on the underside of a dish (**Fig. 3Ai**). After imaging and mapping dot positions, we build an analogue of a ‘cloning ring’ around each dot by jetting surrounding FC40 walls. Instead of jetting circles, we compute the relevant Voronoi diagram – a set of polygons where each contains one dot – and jet polygonal walls around dots (**Fig. 3Aii**; **Fig. S5A**). More fluid can be added to/retrieved from such polygons, within the bounds of advancing and receding contact angles (**Movie 5**). Then, the Poisson limit is ‘beaten’ in the sense that every polygon contains one dot. In practice, some polygons prove unusable – FC40 walls have a thickness (*W*) and two dots may lie <*W* apart (which is too small to accommodate a new wall), and nozzle width may be greater than polygon width (so polygons merge when media is added subsequently). The fraction of such lost polygons inevitably increases as dot density increases; fortunately, densities used for cloning ensure few are lost (e.g., only 2% with 100 dots/cells per 2 cm square; **Fig. 3Aiii**).

**Figure 3.**
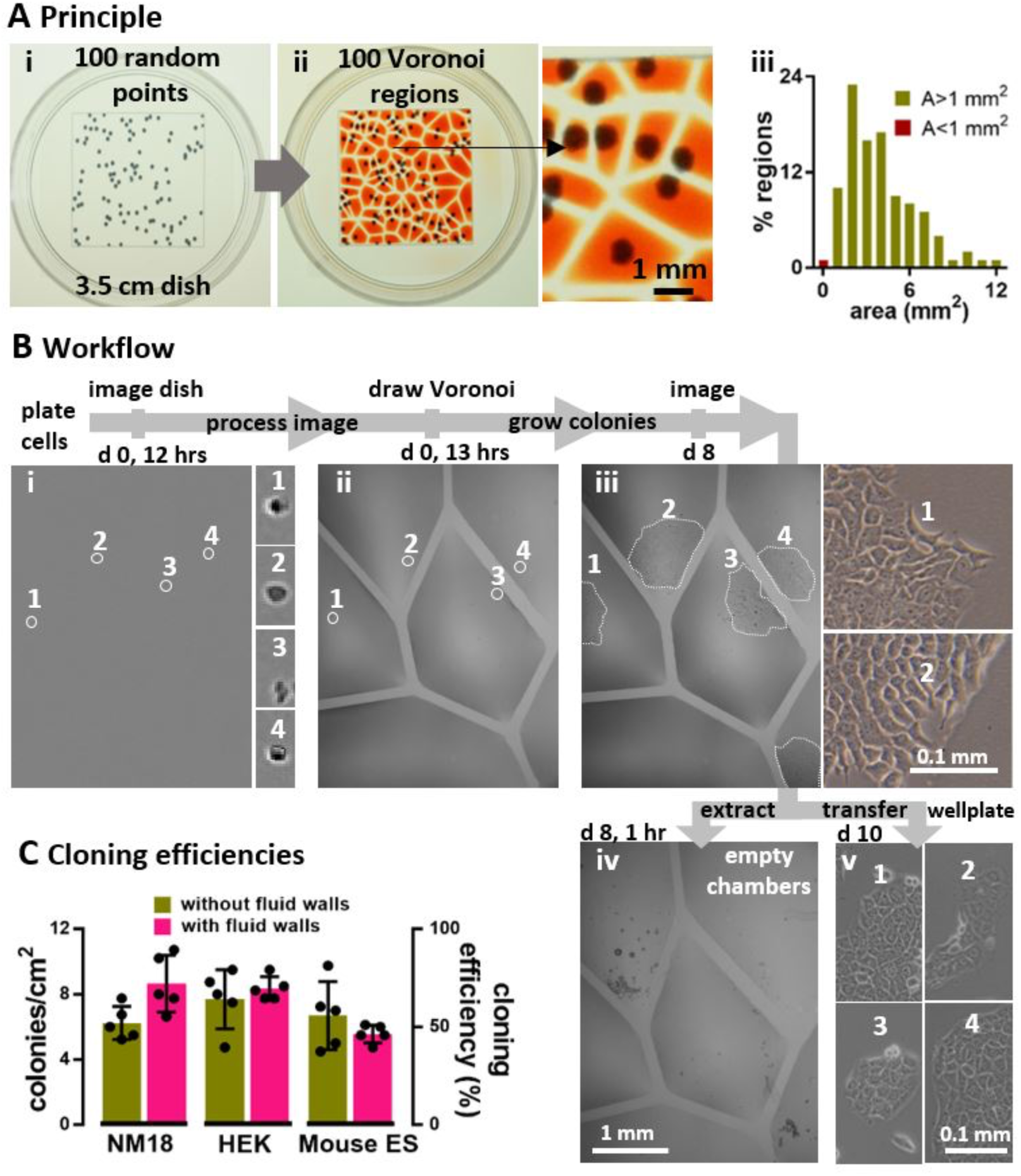
Beating the Poisson limit during cell cloning by limiting dilution. **A.** Principle illustrated using 100 randomly-distributed dots in a 2 cm square printed on clear film placed under a 35 mm dish. (**i**) Dish + dots. (**ii**) After recording dot positions, the Voronoi diagram is determined, polygonal walls built around each dot, and each polygonal chamber filled with dye (zoom shows some chambers). (**iii**) Distribution of areas in this Voronoi diagram. Two polygons had area (A) <1 mm^2^, a threshold determined by the minimum area (hence volume) the infusing needle can access. **B.** Workflow illustrated using NM18 cells (12 cells/cm^2^ or ∼48 in 2 x 2 cm square) plated in a 35 mm dish. Phase-contrast images are of the same region, and numbered zooms show magnifications of different founder cells and their colonies. (**i**) Cells 1-4 12 h after plating. (**ii**) The same cells after jetting surrounding polygons. (**iii**) By day 8, cells develop into colonies (outlined by dotted lines). (**iv**) Emptied polygons (achieved by adding/removing PBS, adding trypsin, incubation, removing most cells). Over-emptying some chambers leaves FC40 (dark blobs) attached to the bottom. (**v**) After removal, each clone was re-seeded in a well in a 12-well plate, and imaged 2 days later. **C.** Cloning efficiencies. Each cell type was seeded (12 cells/cm^2^) in 10 x 35 mm dishes, colonies counted after 8 days, and cloning efficiencies calculated. 5 dishes were used to assess cloning efficiencies conventionally (‘without fluid walls’), and 5 using polygons (‘with fluid walls’). There was no or little significant difference between the two approaches (unpaired t test for HEK, ES, and NM18 gave p = 0.488, 0.3, and 0.033, respectively).

**Figure 3B** illustrates an experiment. Mouse NM18 cells are plated, imaged (**Fig. 3Bi**), positions of viable cells identified, the Voronoi diagram computed, and polygons jetted around each living cell (**Fig. 3Bii**). Once colonies grow (**Fig. 3Biii**), trypsin is added to polygons, cells removed (**Fig. 3Biv**) and transferred to standard 12-well plates, and clones expanded (**Fig. 3Biv**). Cloning efficiencies with 3 mammalian cells (NM18, mouse ES, human embryonic kidney – HEK) are as high as those obtained conventionally (**Fig. 3C, Fig. S5B,C**); moreover, >90% cells identified as viable adherent ones in original images are isolated successfully in polygons, and >95% picked colonies are successfully transferred to (and expanded in) wells in conventional plates. These results show the Poisson limit can be ‘beaten’, with clones picked after 8 days in this case (instead of ≥2 weeks) due to the excellent optical clarity afforded by fluid walls. This approach also gives users confidence that a polygon contains only one cell/colony, so they can forego a 2^nd^ cloning round. More generally, any cell of interested in a complex population can be isolated (e.g., one with a particular morphology or expressing a fluorescent reporter).

**Figure S5.**
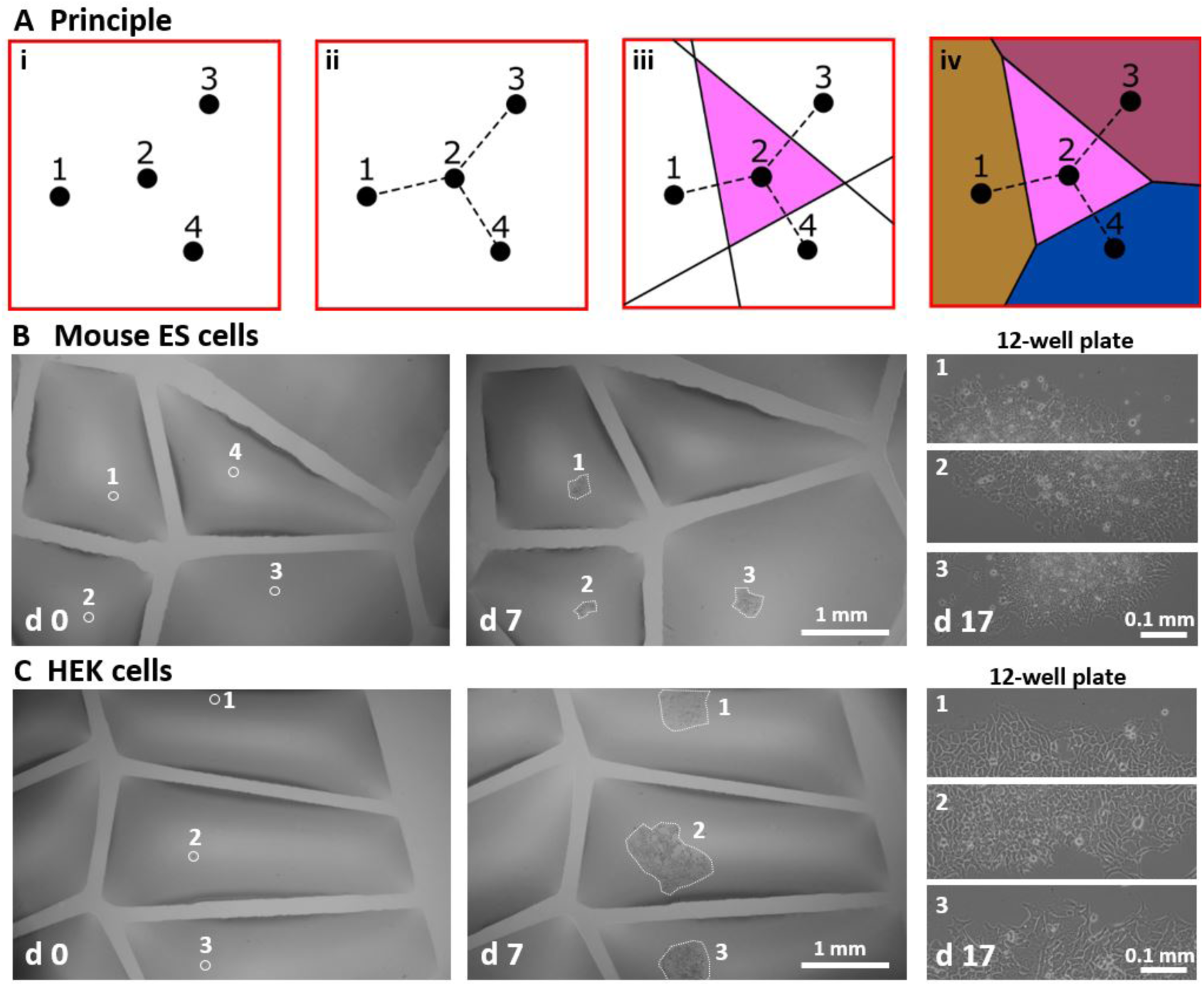
Drawing Voronoi diagrams, and examples of their use. **A.** Principle. (**i**) In this example (4 points randomly-distributed in a square), (**ii**) we draw straight lines between each point (here shown for point 2) and its nearest neighbors. (**iii**) The corresponding Voronoi polygon is formed using half-plane intersections for all lines. (**iv**) The same procedure is applied for other points to produce the Voronoi diagram. **B, C.** Example Voronoi diagrams used during cloning of ES and HEK cells. Details are as Fig. 3, and each pair of images on the left show the same region of the dish immediately after plating cells (d 0; circles surround single cells), and prior to picking colonies (d 7; dotted lines outline colonies). Right-hand panels show views of picked colonies growing in conventional 12-well plates at d 17.

### Sub-culturing cells using contactless jetting

The momentum of an FC40 jet can be insufficient to form a wall but sufficient to dislodge adherent cells from a dish without using trypsin or EDTA (think of a jet-hose cleaning the bottom of a pool). There are two regimes. In one, the nozzle is placed in FC40 above media so jet momentum depresses part of the FC40– media interface to force it down so it plays on attached cells and detaches them (**Fig. 4Ai**). The jet minimizes its area of contact with the aqueous phase when rebounding back to rejoin the overlay (**Movie 6, 7**). Whole colonies can be dislodged, extracted, and replated (**Fig. 4B**). Half a colony can be retrieved, as the other half is kept as back-up (**Fig. 4C, Movie 8**). Naturally, different cell types require different amounts of shear for detachment, with consequential effects on viability. For example, loosely-attached HEKs and NM18 are easily detached as >95% remain viable, whereas tightly-attached ES cells are more difficult to dislodge and ∼5% reattach and regrow (**Fig. S6A**). The jet also produces vortices in chambers that can be used to mix small volumes (e.g., 200 nl human blood; **Movie 9**).

**Figure 4.**
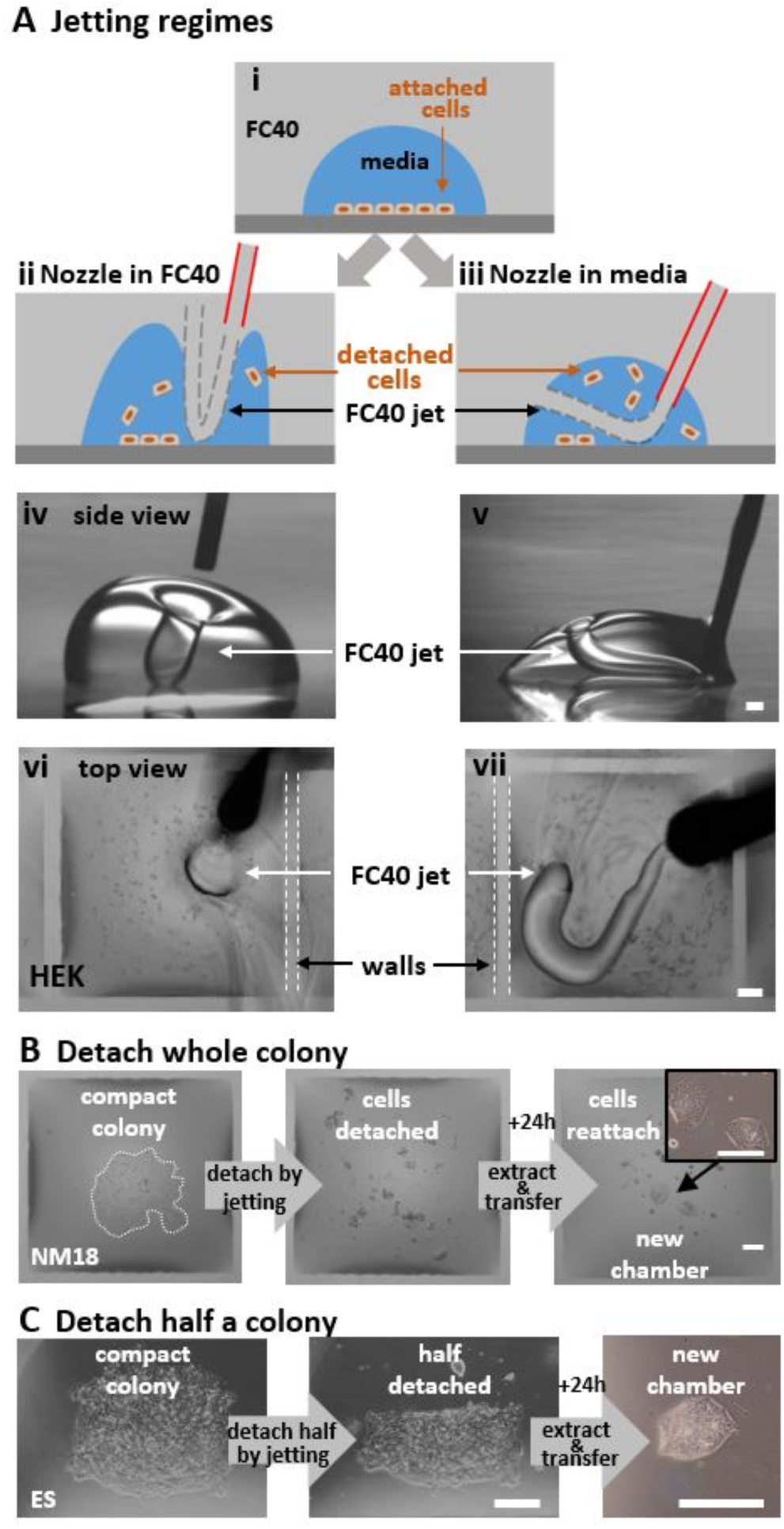
Dislodging attached living cells by jetting. Scale bars: 200 µm. **A.** Two jetting regimes (illustrated using a drop of media and a hand-held jet viewed through a microscope; white dashes outline wall edges). (**i-iii**) With the nozzle in FC40 or in media, a jet can depress the FC40:media interface so it plays on, and detaches, cells. (**iv**-**v**). Side views of the two regimes operating on drops (volumes 3 and 1 µl) with circular footprints (without cells). FC40 jets are invisible. (**vi,vii**) HEKs are plated in a 16 x 16 grid (as **Fig. 1B**), grown for 24 h, and detached using the two regimes and a hand-held nozzle. Remarkably, fluid walls survive. **B,C.** Detaching a whole or half a colony. A colony of NM18 (outlined by dotted white line) or ES cells growing in a chamber (as Fig. 1B) are detached using a nozzle in FC40, detached cells removed and replated in a new chamber, where they grow.

The second regime applies if the hydrophilic nozzle is lowered into media so it is wetted around its circumference. Now, jetted FC40 forms a squirming ‘worm’ surrounded on all sides by media as it makes its way back to the overlay, usually through a ‘hole’ in the chamber ceiling (**Fig. 4A**, right; **Movie 10**). Both regimes allow adherent cells to be sub-cultured without adding enzymes or chemicals, and mix micro-volumes efficiently.

**Figure S6.**
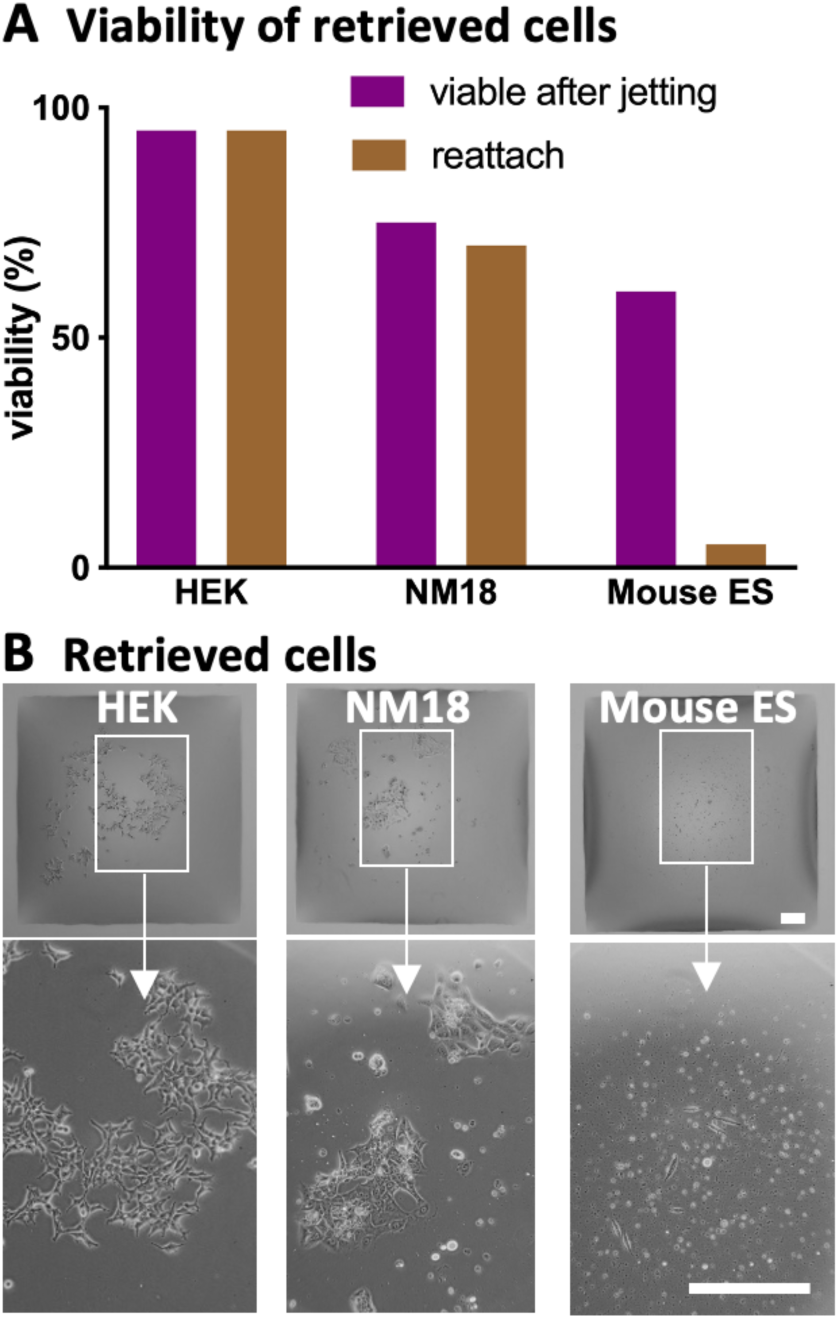
Cell viability after detachment by FC40 jetting. Cells (HEK, NM18, ES) in single colonies in a chamber (as in Fig. 1B) were dislodged by FC40 jetting with the nozzle in FC40 and detached cells retrieved from the chamber; after assessing cell viability by trypan-blue staining, cells were transferred to new chambers and the percentage reattaching assessed. **A.** Viability of retrieved cells. As might be expected, viability and reattachment rates depend on jet momentum required for detachment. Thus, loosely-attached HEKs are easily detached by jetting to remain viable and reattach, whereas tightly-attached ES are detached with difficulty and have poorer viabilities and reattachment rates. **B.** Images of retrieved cells 24 h after replating in new chambers. Bars: 200 µm. Zooms show magnifications.

### A complex perfusion circuit with constant flow

We now illustrate the design, fabrication, and operation of a complex circuit that provides steady flows of fresh media for 7 d as cell grow in an array of chambers. Many conventional circuits doing this have been made (e.g., Khoury *et al*., 2010; Jun *et al*., 2019; Lohasz *et al*., 2019), but they are custom-built and difficult to integrate into workflows in bio-labs. We designed one where every chamber receives fresh media. An external pump drives media through an input conduit (IC), on to one cell chamber (in a 4 x 12 array), and thence to an output conduit (OC), a choke (C), and finally in to the sink (the rest of the dish). The choke acts to minimize pressure differences between input and sink, so equalizing flows through each cell chamber; **Fig. 5Ai**). During tests with red dye, dye intensities in all cell chambers are similar (**Fig. 5Aii**). As fluid walls/ceilings morph during flow to reflect internal pressures, the demonstration that all chambers (which have identical footprints) have similar heights confirms that all experience similar pressures, shear stresses, and flows (**Fig. 5Aiii**; **Fig. S7**). This illustrates another advantage of fluid walls: chamber/conduit height serves as an inbuilt pressure and flow sensor.

**Figure 5.**
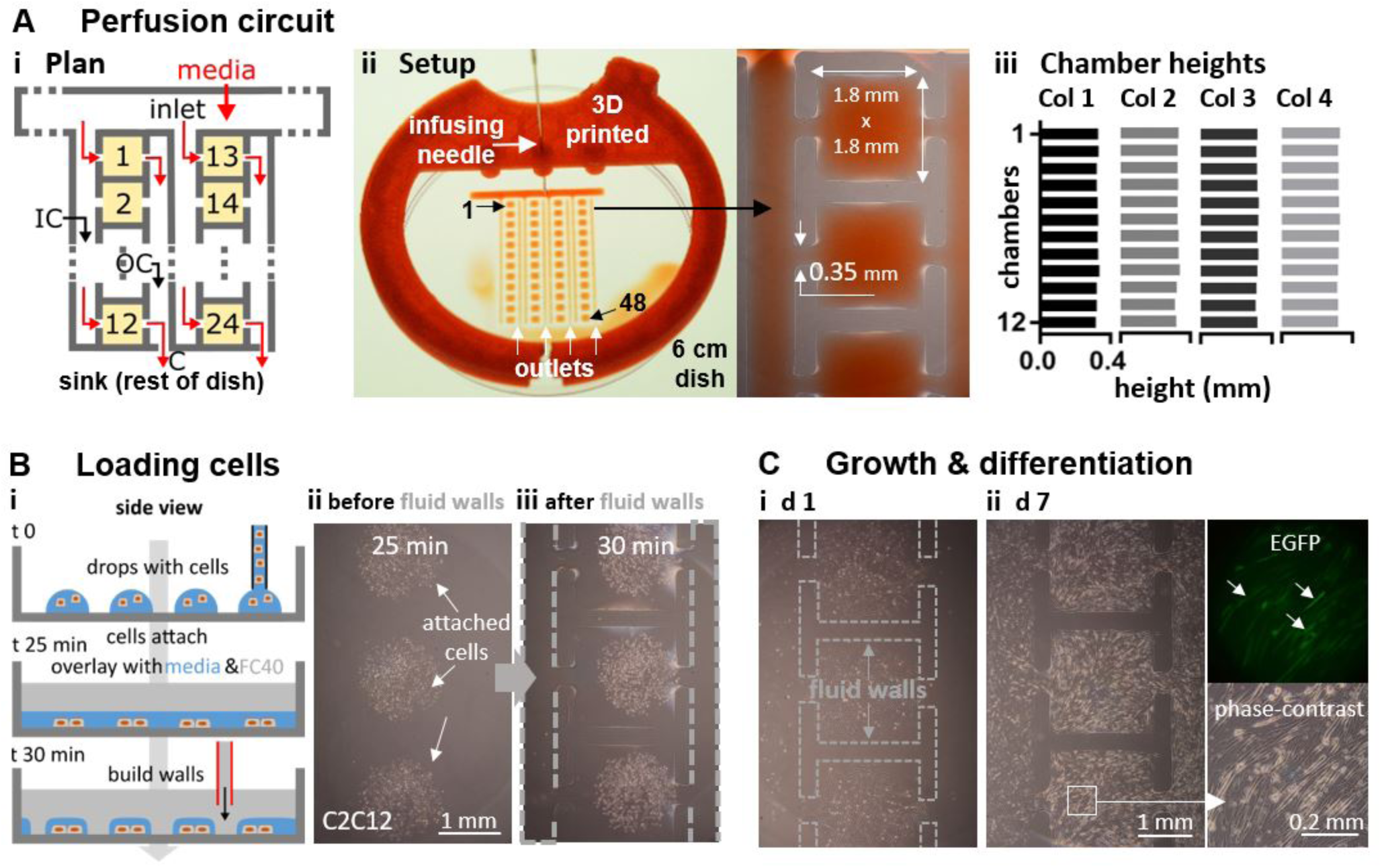
Perfusion circuit for continuous feeding 48 sets of differentiating myoblasts. **A.** (**i**). Circuit plan. Media (± dye) driven by an external pump into the inlet flows (red arrows) through an input conduit (IC), one cell chamber (chambers 1,2,…24 are orange), an output conduit (OC), a choke (C), and out into the dish (the sink). (**ii**) Setup. Red dye is pumped into the inlet through a 3D-printed adapter (red) that fits on a 6 cm dish, through 48 chambers, and out into the dish (zoom shows some chambers). (**iii**) All chambers have similar maximum heights (average ± SD = 325 ± 6.3 µm; assessed using fluorescent dyes as in Fig. S7) showing that all experience similar pressures, shear stresses, and flows. **B.** Growth and differentiation of mouse C2C12 myoblasts. (**i**) Workflow. Myoblasts are deposited in drops, the dish filled with media and FC40 added, and finally the circuit is jetted around them. (**ii,iii**) Cells before/after building fluid walls. Dashed lines: edges of some walls. **C.** Images of chambers as media replenished (1 µl/day/chamber); myoblasts grow to fill chambers, form syncytia, and express EGFP-DOK7 (day 7). Fluorescence and phase-contrast zooms: arrows mark fluorescing syncytia each containing >20 nuclei with length >200 µm – indicative of differentiation into myotubes.

Circuit operation is demonstrated using mouse C2C12 myoblasts that can differentiate over 7 d into mature myotubes; they fuse to form syncytia and express components of the neuromuscular junction – including the adaptor protein, DOK7 – which in this case is tagged with EGFP (Cossins *et al.*, 2012). The undifferentiated (non-fluorescent) myoblasts can be added to each chamber immediately after fabrication; then, as chambers have Laplace pressures lower than flanking conduits, deposited cells remain in chambers. While this method is quick, it has the drawback that cells are exposed to equilibrating flows (and shear stresses) as they settle, and this can adversely affect some cell types (not shown). Therefore, cells are deposited as drops (in air) on a virgin dish at positions corresponding to those where the 48 chambers will eventually be built; on incubation, cells attach under shear-free conditions (**Fig. 5Bi**). Next, media is added to the whole dish to merge drops, FC40 overlaid, and the circuit built around the 48 groups of living cells (**Fig. 5Bii, iii**). The dish is now placed in a CO_2_ incubator, and fresh media slowly pumped into the circuit over 7 d. Then, cells grow and differentiate into myotubes, form syncytia, and become fluorescent (**Fig. 5C**; **Fig. S7D**). This shows that flows through a complex circuit can be equalized by judicious design, and that cells flourish and differentiate normally within it. Although growth increases numbers and some cells migrate towards inlet and outlet channels to eventually meet those from neighboring chambers, microscopic analysis is easily restricted to those cells that remain in any particular chamber. If required, any chamber can be disconnected from the rest of the circuit by building new FC40 walls across input and output conduits, and its contents extracted for downstream analysis.

**Figure S7.**
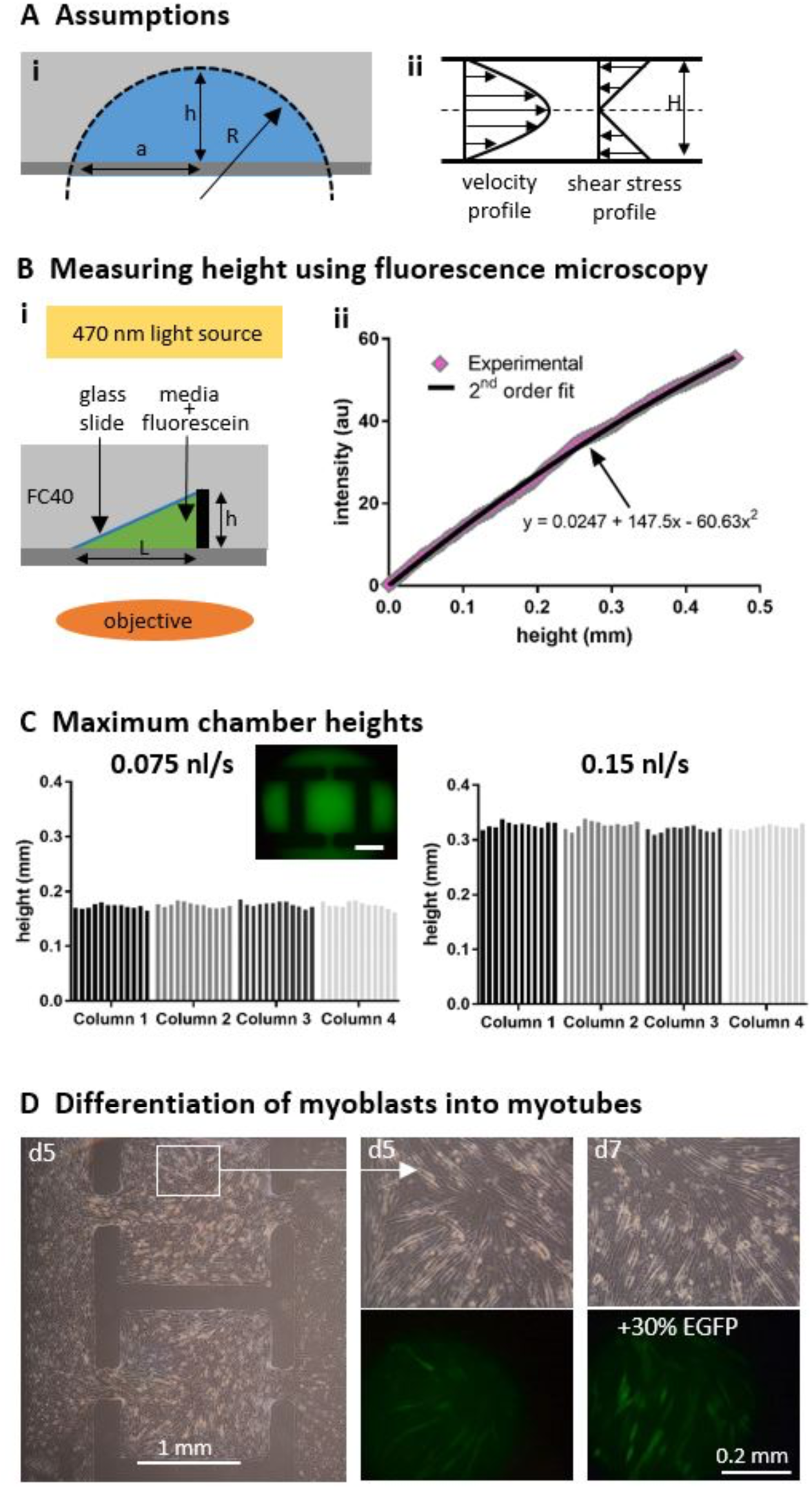
Confirming that cells in different chambers in the perfusion circuit in Figure 5 experience the same pressures and velocity profiles (and so rates of media replenishment and shear stresses). **A.** Assumptions. **(i**) The 3D geometry of a cell chamber is complex, resembling that of a spherical cap sitting on a rectangular base (Soitu *et al*., 2018); for simplicity, we assume it is a perfect cap of a sphere (2a = chamber width, R = sphere radius). (**ii**) Then, the velocity and shear-stress profiles between bottom and top of any section through a chamber are functions of height, H, at that point in the section. Identical heights in all chambers indicate that cells in each chamber experience similar pressures, shear stresses, and flows. **B.** Measuring maximum chamber height using fluorescence microscopy. (**i**) Construction of a calibration curve relating fluorescence intensity to height of fluorescein. A mixture of cell media and fluorescein is added between a glass slide propped against a pillar (height, h) and the bottom of a Petri dish, FC40 overlaid, the construct imaged, and the intensity at each point along the slide recorded. (**ii**). Relation between fluorescence intensity (arbitrary units, au) and height (calculated knowing dimensions of the triangle in (i)). Black line: best fit of a second-order polynomial. **C.** Maximum chamber heights in circuits perfused at different rates. Media plus fluorescein was perfused through chambers in circuits (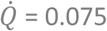and 0.15 nl/s; data for 0.15 nl/s is reproduced from Fig. 5Aiii), fluorescence images collected (inset: fluorescence image of part of a circuit, bar = 1 mm), and chamber heights calculated using the calibration curve. Average heights were 175 µm ± 5 µm and 325 µm ± 6.3 µm SD at 0.075 and 0.15 nl/s, respectively. This indicates that for a given circuit and flow rate, heights of different chambers are similar, and that chamber height changes with flow rate (as fluid walls morph above unchanging footprints to accommodate the pressure difference). As chamber heights reflect both 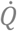 and velocity profiles, each chamber in a circuit experiences the same rate of media replenishment and shear stress. **D.** Control test to show the increase of EGFP signal with differentiation. C2C12 cells were cultured for 7 d in the perfusion circuit. Insets show phase-contrast and fluorescence images of the same location inside a chamber at d 5 and d 7. For an exposure time of 1 s, the average intensity of signal per pixel increases by 30% between the two time points, indicating increased expression of the tagged DOK7 (and so differentiation into a myotube).

## DISCUSSION

We describe a general and rapid method to fabricate microfluidic circuits on a standard polystyrene Petri dish. A micro-jet of FC40 is projected through FC40 and media on to the bottom of a dish; as FC40 hits the dish, it washes away media to leave a liquid wall of FC40 tightly ‘pinned’ to the substrate by interfacial forces (**Fig. 1A**). Circuits with almost any imaginable 2D shape can be built in seconds to minutes (**Fig. 1B-H**). We establish conditions required to build stable fluid walls (**Fig. 2**), drive flow through conduits without using external pumps (**Fig. S3**), and create ‘valves’ that can be opened and closed at will (**Fig. S4**). We develop an approach for cloning dilute suspensions of mammalian cells that can beat the Poisson limit in the sense that almost every cell in the initial population is confined within its own fluid wall (**Fig. 3**). We extend the approach to sub-culture attached cells without adding trypsin or EDTA (**Fig. 4**), and to feed arrays of mammalian cells over long periods (**Fig. 5**). Unfortunately, evaporation from static volumes of less than ∼1 nl inevitably limits construction of ever-smaller devices (even though the solubility of water in FC40 is <7 ppm by weight).

In summary, we have developed a contactless method for fabricating microfluidic devices on standard Petri dishes in minutes. As it uses materials familiar to biologist, we anticipate it will find wide application in biomedicine.

## MATERIALS AND METHODS

### General reagents

All reagents and materials were purchased from Sigma Aldrich (St. Louis, Missouri), unless otherwise stated. FC40^STAR^ (iotaSciences Ltd, Oxfordshire, UK) is FC40 treated using a proprietary method that improves wall formation by jetting, and throughout we use the term ‘FC40’ to refer to it. Water-soluble dyes (e.g., Allura Red, toluidine blue, resazurin) are used where indicated.

### Cells, cell culture

Whole human blood (anti-coagulated with EDTA) was obtained from anonymous blood donors through the National Health Service (NHS) Blood and Transplant Service (Oxford, UK); ethical approval was provided by The Interdivisional Research Ethics Committee of Oxford University (R63966/RE001).

Adherent human embryonic kidney cells (HEKs) were grown in DMEM (Gibco, Gaithersburg, MD) + 10% FBS; these HEKs were genetically-modified reporter cells (NF-κB/293/GFP-Luc™ Transcriptional Reporter Cell Line; System Biosciences, catalogue number TR860A-I), but reporter activity is not relevant here.

Mouse embryonic stem (ES) cells (EK.CCE line, derived from a single XY blastocyst-stage embryo of strain 129/Sv/Ec; Robertson *et al*., 1986) were routinely cultured in DMEM supplemented with 15% FBS, 1% penicillin/streptomycin (Gibco, #15140122), 0.1 mM 2-mercaptoethanol, 1% glutamine (Invitrogen, Loughborough, UK), 1% minimum essential media nonessential amino acids (Gibco), 1 mM sodium pyruvate, and 1000 U/mL leukemia inhibitory factor (LIF, ESGRO; Merck, Darmstadt, Germany; #L5158) on gelatin-coated plates (Merck, #ES-006-B). Such plates were used for **Figures 3C** and **S5B**.

Mouse mammary tumor cells (NM18, a derivative of the NMuMG line; Deckers *et al*., 2006) were cultured in DMEM supplemented with 10% FBS, 1% penicillin/streptomycin, and 0.1% insulin (Sigma Aldrich, #10516).

C2C12, an immortalized mouse myoblast (Cossins *et al*., 2012), were cultured in DMEM + 15% FBS. For the perfusion circuit in **Figure 5**, cells were seeded in DMEM + 15% FBS, and the FBS was reduced to 6% after 4 d, and to 2% after 5 d.

### Printing and operating circuits

All circuits were made on tissue-culture-treated surfaces – 60 or 35 mm circular dishes (Corning, Merck product 430166 or #430165) or rectangular uniwell plates with overall dimensions of 96-well plates (Thermo Fisher Scientific, Waltham, MA) using custom-written software (see also below) and modified ‘Freestyle’ or ‘Pro’ printers (iotaSciences Ltd, Oxfordshire, UK). Each printer consists of a 3-axis traverse, which controls movement of stainless steel needles (Adhesive Dispensing Ltd, Milton Keynes, UK) – usually a 60 µm inner-diameter nozzle jetting FC40 (outer diameter 0.5 mm), and a 0.5 mm outer-diameter needle (inner diameter 0.25 mm) used to add/remove media to/from chambers. The two needles are each connected via Teflon tubing to a programmable syringe pump. The Freestyle accepts 60 and 35 mm dishes, while the Pro has a larger working area and less mechanical backlash, so it was used to print the circuits in **Figures 1D, 1E, 1G, S1** and **5A**. Flows through circuits were driven by programmable syringe pumps (Harvard PhD Ultra I/W) connected via Teflon tubing to blunt stainless-steel dispensing needles (outer diameter 0.5 mm); needles were lowered through FC40 into the circuit at the appropriate position, when fluid walls spontaneously self-seal around inserted needles.

DMEM plus 10% FBS was used to make all grids and circuits; when the term ‘medium’ is used, it should be assumed that 10% FBS is also present. Routinely, to prepare a grid or circuit, 1 ml medium is pipetted into a 60 mm dish to wet the entire bottom surface and ∼0.9 ml retrieved to leave a thin film. This is then overlaid with 2-3 ml FC40 and the dish placed on the printer. Standard conditions used for jetting FC40 when making most circuits were: *D*_*nozzle*_ = 60 µm, *H* = 0.4 mm, 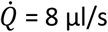, *V*_*traverse*_ = 10 mm/s. A tightly-fitting 3D-printed circular sleeve around each dish acts as a positioning ring to ensure the dish can be removed and added back on to the printer in the original orientation. The adapter in **Fig. 5Aii** used to hold the needle feeding the perfusion circuit in a 60 mm dish was 3-D printed with polylactic acid (rigid.ink, Wetherby, UK).

Patterns in **Figures 1C, 1D, 1E, 1G, 1H, S1**, and **5A** were constructed after reproducing the desired circuit in Inkscape (inkscape.org) and conversion to G-code, the programming language used by the printer. Patterns for **Figures 1D, E, G** and **1H** were obtained from Wikipedia, CanStockPhoto, http://www.udel.edu and WikiArt, respectively.

For **Figures 3** and **S5**, ImageJ (Rasband, 1997-2018) was used to detect and highlight attached single cells. Images were then processed in MATLAB (MathWorks, Cambridge, UK), where coordinates of single cells were recorded and Voronoi diagrams subsequently generated. The code used is available in Supplementary Materials. Finally, Inkscape was used to convert the diagrams into the G-code used to print polygons. Areas and centroids of each polygon were also computed, and the volumes to be added to each polygon calculated using the formula V_cap_ = (A_wetted_ /3.7)^3/2^ – where V_cap_ is the maximum volume that can be added to a polygon assuming an advancing contact angle of 50° (Walsh *et al.*, 2017), and A_wetted_ is the area of the polygon. The same volumes were removed from, and added to, each polygon when refreshing media, and during colony retrieval (which involved emptying chambers, PBS addition and removal, trypsin addition, and removal of the resulting cell suspension). Retrieved cells were plated in 12-well plates (Greiner Bio-One, Kremsmünster, Austria; #665180) and allowed to reattach and grow.

### Imaging

Images of dishes in **Figures 1B, 1D, 1E, 1G, 1H, S1, S2C, 3A, S3C, 4A, S4C, 5A**, and **S7A** were taken using a digital SLR camera (Nikon D610). All insets in these figures, and fluorescence images in **Figures 5C** and **S7D** were collected using a zoom lens and digital SLR camera (Nikon D7100 DSLR) connected to an epi-fluorescent microscope (Olympus IX53; 1.25X, 4X, 10X, 25X objectives) with translation stage and overhead illuminator (Olympus IX3 with filters).

For **Figure 3**, 35 mm dishes were imaged using a phase-contrast live-cell imaging system and a 4x objective (IncuCyte Zoom, Sartorius, Gottingen, Germany).

For fluorescent images of cells in **Fig 5C** and **Fig S7D** an exposure time of 1 s was used. Quantification of EGFP intensity was done in ImageJ by subtracting the background and averaging the intensity across the entire image. Brightness was then increased by 40% for these images for a better visualisation of cells expressing EGFP.

**Movies 3, 6, 7**, and **10** were taken using a pendant-drop instrument (First Ten Angstroms 1000, Cambridge, UK).

### Statistical analyses

Statistical analyses were performed using GraphPad Prism (San Diego, CA).

## ACKNOWLEDGEMENTS

We thank the labs of David Beeson (for C2C12 cells), Elizabeth Robertson (for mouse ES cells), and David Greaves (for access to the IncuCyte), and iotaSciences Ltd for providing printers and FC40^STAR^.

This work was supported by iotaSciences Ltd (who provided scholarships for C.S., C.D.), the William H.G. FitzGerald Scholarship (N.S.-K.), and a Royal Society University Research Fellowship (A.A.C.-P.), the Impact Acceleration Account of the Biotechnology and Biological Sciences Research Council (P.R.C. and E.J.W.), and awards from the Medical Research Council under the Confidence in Concept scheme (MC_PC_15029 to P.R.C. and E.J.W).

## AUTHOR CONTRIBUTIONS

C.S., P.R.C., and E.J.W. designed research; C.S., N. S-K., C.D., and E.J.W. performed experiments; all authors wrote the paper.

## SUPPLEMENTARY MOVIES

**Movie 1.** Jetting a grid. The movie runs at 4x speed. A Petri dish (6 cm) – filled with a skim of media mixed with blue dye plus an FC40 overlay – sits on a printer, and the nozzle of a hollow needle (*D*_*nozzle*_ = 60 µm) is held by a 3-way traverse in FC40 above media in the dish (*H*= 0.4 mm). The movie begins as the pump infuses FC40 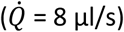 and the traverse traces the path of a grid (*V*_*traverse*_ = 10 mm/s). The invisible jet of FC40 pierces the layer of media, pushing it aside to contact the substrate. As FC40 wets polystyrene preferentially, it remains firmly pinned to it. A grid with 256 aqueous chambers, each surrounded by transparent FC40 walls, is created in <2 min.

**Movie 2.** A 16 x 16 grid withstands violent agitation. The movie runs in real time. A grid like the one in Figure 1B was constructed, each chamber filled with 600 nl red dye, and the dish placed on a shaker.

**Movie 3.** Jetting Novec 649 from a nozzle submerged in FC40 confirms that jet width increases with distance from the nozzle. The movie runs in real time (10 frames/sec). This side view depicts a nozzle of a 34G needle (*D*_*nozzle*_ = 80 µm, outer diameter 220 µm) submerged in FC40. Then, another fluorocarbon (i.e., Novec 649) – chosen because it is miscible with FC40 but has a different refractive index so interfaces between the two transparent liquids can be seen – is jetted from the nozzle (*D*_*nozzle*_ = 60 µm) through FC40. Jet width increases with distance from the nozzle (see also Deshpande and Vaishnav, 1982). Turbulence at the end of the movie results from Novec 649 rebounding from the substrate.

**Movie 4.** A microfluidic valve. The movie plays at 1x or 4x real time as indicated. It illustrates the experiment shown in Figure S4. The 6 cm dish initially contains 5 identical circuits, each a reservoir connected to a conduit 1 mm wide with a dead-end, and each bounded by one continuous wall. Then, the printer adds 2 µl red dye to each reservoir (walls are intact and dye stays within each circuit), jets media to open end walls (red dye spills out into the dish), jets FC40 to rebuild end walls, adds 2 µl blue dye to reservoirs to test for leaks (no blue dye escapes), and jets media to reopen end walls (so now blue dye does escape).

**Movie 5.** Visualizing a Voronoi diagram. The movie is speeded up 4x, and illustrates the kind of experiment shown in Figure 3A where an area of 20 x 20 mm in a 35 mm dish was subdivided into 64 Voronoi polygons, each containing one dot. Polygons are initially invisible due to the low volume each contains. The movie shows addition of media + black dye by a printer through FC40 to each polygon (added volume is proportional to polygon area, and ranges from 0.25–2.8 µl).

**Movie 6.** FC40 from a nozzle held above drops of media deforms FC40–media interfaces so a depressed section plays on the bottom. The movie runs in real time (10 frames/sec). It depicts a side view of a nozzle (a 34G needle, *D*_*nozzle*_ = 80 µm, outer diameter 220 µm) held below the surface of FC40 but above drops of media (shaped like caps of spheres with volumes of 1–3 µl) sitting in a dish. The movie begins as the nozzle jets FC40, and traverses above drops. Although the jet is invisible, it deforms sections of the FC40–media interface in each drop as it passes. These sections are forced down to the substrate, and can be used to detach adherent cells.

**Movie 7.** Jetting Novec 649 from a nozzle held above a drop of media deforms the FC40–media interface so a depressed section plays on the bottom. The movie runs in real time (10 frames/sec). In Movie S3, FC40 is invisible, and there is no indication of any turbulent flow after the jet leaves the drop. Here, the jet contains another fluorocarbon (Novec 649) that is miscible with FC40 but has a different refractive index so interfaces between all 3 liquids can be seen. Initially the nozzle (34G needle, *D*_*nozzle*_ = 80 µm, outer diameter 220 µm) is held below the surface of FC40 but above a 1 µl drop of media. Novec 649 is then jetted on to the drop, the nozzle traverses back and forth above the drop, and a depressed section of the FC40–media interface plays on the bottom.

**Movie 8.** Detaching half a stem-cell colony growing in a chamber in a grid. The movie runs in real time, and images were collected on a microscope from above. The movie depicts removing part of a living colony of mouse ES cells from a chamber (as in Fig. 1B) using a hand-held FC40 jet (*D*_*nozzle*_ = 60 µm; 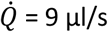). The nozzle is positioned in FC40 and above the media–FC40 interface throughout, and the jet depresses a section of the interface so it plays on the bottom of the chamber to detach some cells in the colony. Figure 4C illustrates accurate removal of half a colony using the printer, but then imaging through a microscope from above (as in this movie) is impossible.

**Movie 9.** Mixing blood using the regime described in Figure 4A where the nozzle is held below an FC40:aqueous interface. The movie runs in real time. A grid like the one in Figure 1B was made, 0.2 µl whole human blood added to two chambers, and a manually-held jet used to create vortices in chambers, and so mix the blood.

**Movie 10**. Jetting FC40 from a hydrophilic nozzle submerged in a drop of media yields a ‘squirming worm’ (which can be used to detach cells). The movie runs at 5x speed (10 frames/s). This side view depicts a 1 µl drop of media (initially shaped like the cap of a sphere) under FC40, and the hydrophilic nozzle (*D*_*nozzle*_ = 80 µm) is below the surface of FC40 but above the drop. After 1s, the pump jets FC40 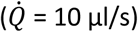 on to the drop to deform the FC40–media interface. After 5 s, the hydrophilic nozzle is lowered towards the drop. By 12 s, the nozzle enters the drop and is wetted around its circumference; consequently, the FC40 jet is surrounded on all sides by media as it leaves the drop to rejoin the overlay (which it does at the back, and so out of sight). After ∼24 s, flow rate is increased 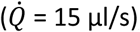, and the jet rearranges and extends from the nozzle to the right to appear as a ‘squirming worm’. By 35 s, the ‘exit hole’ of the jet from the drop is clearly visible. At ∼85 s, the flow rate is reduced 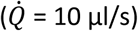, and at ∼125 s increased again 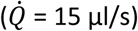. The worm continues to squirm in the drop until the end of the movie.

### Code used to create Voronoi diagrams

~~~
%read image with single cells to be analysed.
image = imread(‘single_cells.tif’);
%get the size of the image
[rows, columns, numberOfColorChannels] = size(image);
%show the image
imshow(secondlevelTIFF);
%project a grid on top of the image to help with the identification of single cells
grid_lines_every = 1000;
for row = 1 : grid_lines_every : rows
 line([1, columns], [row, row], ‘Color’, ‘r’);
end
for col = 1 : grid_lines_every : columns
 line([col, col], [1, rows], ‘Color’, ‘r’);
end
%record position of each cell
X = []; Y = []; points = []; i = 0; datacursormode on;
while (i < 1000)
 dcm_obj = datacursormode(figure(1));
 pause
 cursor_info = getCursorInfo(dcm_obj); position = cursor_info.Position;
 X = [X, position(1)]; Y = [Y, rows - position(2)]; i = i + 1;
end
datacursormode off
%store positions in txt files
dlmwrite(‘Xpos.txt’, X); dlmwrite(‘Ypos.txt’, Y);
%apply Voronoi diagram to the set of points
voronoi(X,Y);
~~~

